# Dynamic live/apoptotic cell assay using phase-contrast imaging and deep learning

**DOI:** 10.1101/2022.07.18.500422

**Authors:** Zofia Korczak, Jesús Pineda, Saga Helgadottir, Benjamin Midtvedt, Mattias Goksör, Giovanni Volpe, Caroline B. Adiels

**Affiliations:** Department of Physics, University of Gothenburg, Sweden

## Abstract

Chemical live/dead assay has a long history of providing information about the viability of cells cultured in vitro. The standard methods rely on imaging chemically-stained cells using fluorescence microscopy and further analysis of the obtained images to retrieve the proportion of living cells in the sample. However, such a technique is not only time-consuming but also invasive. Due to the toxicity of chemical dyes, once a sample is stained, it is discarded, meaning that longitudinal studies are impossible using this approach. Further, information about when cells start programmed cell death (apoptosis) is more relevant for dynamic studies. Here, we present an alternative method where cell images from phase-contrast time-lapse microscopy are virtually-stained using deep learning. In this study, human endothelial cells are stained live or apoptotic and subsequently counted using the self-supervised single-shot deep-learning technique (LodeSTAR). Our approach is less labour-intensive than traditional chemical staining procedures and provides dynamic live/apoptotic cell ratios from a continuous cell population with minimal impact. Further, it can be used to extract data from dense cell samples, where manual counting is unfeasible.

---

Evaluating cellular behaviour in vitro is essential when assessing their responses to external influences of physical and chemical nature. One standard evaluation method is to use a live/dead assay. Such an assay involves staining the cells with fluorescent dyes followed by fluorescence microscopy imaging and subsequent cell counting (live or dead). Unfortunately, this method comes with many disadvantages. Chemical staining and fluorescence imaging procedures can disrupt or harm the cells, [1, 2] and the fluorescent dyes themselves can even be cytotoxic.[3–5] Moreover, even if performed on live cells, standard staining protocols require dissolving chemical dyes in buffers where the cells do not receive the necessary nutrients. This may harm them or, at the very least, affect the equilibrium of the sample. Therefore, chemical staining is performed as an endpoint assay, sacrificing samples at every chosen time-point of the experiment. If viability information is required during the experiment, it is necessary to prepare enough cell cultures that may be sacrificed during the course of the experiment, making it impossible to carry out longitudinal experiments on the same cell population.

Standard in vitro cell culturing is performed in static conditions, where control of the external environment is limited, and dynamic time-lapse experiments are challenging to achieve. Further, due to the far from in vivo-like environment, statically cultured cells may not display their inherent morphology, functionality or viability. In contrast, microfluidics and organs-on-chips resemble such environments and can be tailored for different cell conditions. [6–8] Such platforms offer high control of the extracellular environment and, when optically transparent, enable monitoring of dynamic cell responses over time. However, microfluidics also suffers from the drawbacks of standard viability assays, as it also requires multiple devices that can be sacrificed throughout the experiment (Fig.1a). Consequently, longitudinal viability data collection is time-consuming and labour-intensive, and dynamic cell behaviour observations on the same population of cells over time are still impossible.

**FIG. 1.**
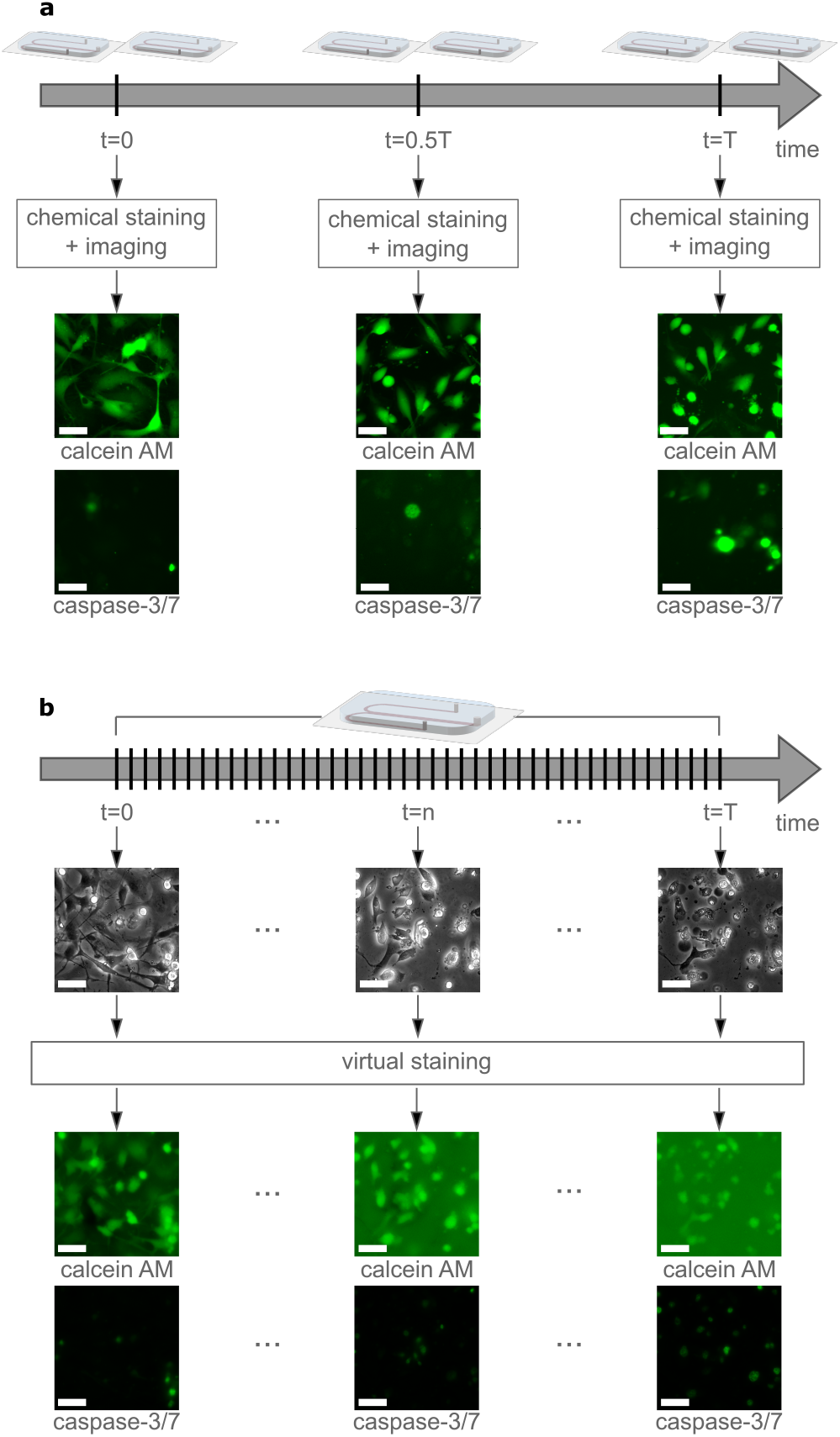
Retrieving cell viability information from inside a microfluidic device. (a) The standard procedure consists of performing chemical staining on-chip, sacrificing two devices at each intended time point, e.g. beginning, half-time and the end of a long experiment (days) and imaging them with a fluorescence microscope. This approach is time-consuming and workload intensive only providing snapshots of the cellular behaviour. Scale bar is 50 µm. (b) The developed method based on deep learning uses two neural networks, one for generating virtually-stained images from phase-contrast images and the other to count cells on previously generated images. The method allows for probe-free, non-invasive and dynamic longitudinal cell response evaluation on the same cell population. Scale bar is 50 µm.

Another combination of viability dyes is required when applying rapid environmental changes using microfluidics. Instead of staining already dead cells, it is advantageous to identify unhealthy cells as early as possible in the apoptotic process. If the apoptotic process is observed early, the extracellular environment can be changed, with the possibility to reverse the active process of programmed cell death.[9] However, also apoptotic dyes suffer from the same limitations previously mentioned.

Thanks to recent advances in deep learning, several of the abovementioned limitations can be overcome. Virtual staining of subcellular structures like cytoplasm or nuclei, or H&E analysis of tissues has been achieved using bright-field[10] or quantitative phase microscopy[11–13] images as input. This approach allows users to retrieve quantitative information from their samples without affecting cells’ environment by exposing them to harmful chemical dyes or fluorescent illumination.

Here, we present an analysis method for detecting and discriminating live or apoptotic cells using deep learning. Detecting apoptotic rather than dead cells enables dynamic and real-time perturbation studies. We propose a conditional generative adversarial neural network (cGAN)-based method using phase-contrast images to generate virtually-stained live or apoptotic cells. Subsequently, we apply a self-supervised single-shot deeplearning technique (LodeSTAR) to count the cells in the stained images.[14] Our method can provide long-term cell viability data on the same population throughout the whole experiment, without the administration of chemical dyes (Fig.1b).

In this study, we cultured human endothelial cells under constant perfusion in a custom-made microfluidic device. In vivo, endothelial cells are constantly exposed to shear stress via perfusion 17] and optimally should be cultured alike in vitro. We used a microfluidic device designed as a universal platform for dynamic mammalian cell studies of either shear stress or gradient exposure. The device was fabricated in polydimethylsiloxane (PDMS) using a master manufactured via soft lithography as a template.[18, 19] It consists of a 1 mm wide centre channel and two 500 µm side channels with a height of 49 µm (a schematic of the device can be seen in Fig.1 and in Fig.S1). Before the experiments, the devices were precoated with 1:100 v/v collagen solution in sterile water (collagen type I from rat tail, Sigma-Aldrich) to create a suitable substrate for cell attachment and growth inside the microchannels.

We used two different chemical dyes to assess the cells’ viability. To account for live cells, we used calcein AM (Thermo Fisher), a lipid-soluble esterase substrate that can passively cross the cell membrane. Inside live, correctly functioning cells, it is converted by intracellular esterases into a green fluorescent molecule (calcein). Even if calcein is considered to remain inside the cells as long as the cell membrane is intact, [20] it suffers from a slow spontaneous leakage.[21] Previous non-steady-state studies have also shown that calcein can self-quench when the dye reaches high enough concentrations.[22, 23] Yet another limitation of this livecell dye is that hydrolysed (i.e. fluorescent) calcein molecules can remain inside the cells’ cytoplasm even though the apoptotic metabolic process has been initiated. [24–26] To account for apoptotic cells, we used CellEvent™ Caspase-3/7 Green Detection Reagent (in short, caspase-3/7), also from Thermo Fisher, that initially is a non-fluorescent conjugate. In early apoptotic cells, it binds to DNA thanks to caspase metabolic processes activation and emits a green fluorescent signal. A minor drawback of this dye, reported by the manufacturer, is that also healthy cells display a minimal positive caspase signal. Because of the spectral overlap between these two dyes, two different chemical staining solutions were prepared using the endothelial cell complete medium with the final concentration of 2 µM for calcein AM and 10 µM for caspase-3/7.

To monitor cells’ transition from living to apoptotic over time, we kept cells chemically-stained at all times to follow this process. Human dermal microvascular endothelial cells, HMEC-1 (ATCC^®^ CRL-3243™) were maintained in a static environment at 37°C with 5% CO_2_ according to the manufacturer’s protocol. On the day of the experiment, cells were detached, centrifuged at 100 rcf (relative centrifugal force) for 7 min and seeded (approximate density of 3.4 * 10^7^ cells/mL) into the coated microfluidic devices. Cells were left to sediment for 2 hours at 37°C to attach to the collagen coating and acquire their typical elongated morphology. We used two different inflow velocities, 0.1 µL/min and 2 µL/min, anticipating that it would result in distinct cell behaviours, i.e., display different dynamic live/apoptotic ratios. Endothelial cells are thin (below 10 µm thickness[27]), so we estimate that the microfluidic-hosted HMEC-1 cells would experience shear stress values of 4 mPa and 80 mPa for low and high flow, respectively. These values lie within the physiological range of shear stress values found in blood capillaries.[28]

The microfluidic device was placed in an on-stage incubator (Chamlide TC, Live Cell Instrument) and cells were monitored using an inverted epi-fluorescence microscope (DMI6000B, Leica Microsystem) with 20x magnification (HCX PL FLUOTAR, NA 0.40, Leica Microsystems). The automated time-lapse imaging started jointly with the constant infusion of a respective staining medium. Sequential imaging with phase-contrast and fluorescence microscopy continued for up to 12 hours.

From the time-lapse phase-contrast images, a cGAN was trained to generate virtually-stained fluorescence images of live (simulating calcein AM) and apoptotic (simulating caspase-3/7) cells. The data from the low flow experiments were used for the cGAN training and internal validation. Because of the spectral overlap of the two dyes, separate data sets were acquired for each of them. The training data comprised 1440 pairs of phase contrast and fluorescence images for live cells and 1980 pairs for apoptotic cells. The internal validation data set consisted of 450 pairs of images for live cells and 792 for apoptotic cells. We used the high-flow data as external validation data to evaluate the network performance (1170 pairs of images for live cells and 3668 for apoptotic cells).

Our cGAN consists of three neural networks (see Fig.S2). First, a generator receives phase-contrast images as input and generates virtually-stained fluorescence images of live and apoptotic cells. The second network is a discriminator, which aims to distinguish the generated images from the images of chemically-stained cells (authentic images), classifying them accordingly. And the third one is a perceptual discriminator that exploits the perceived similarities between the generated and authentic images. As we train these three networks simultaneously, the generator gradually becomes more adept at generating virtually-stained images that fool the discriminator. The discriminator, in turn, becomes better at distinguishing images of chemically-stained cells. The perceptual discriminator is novel in virtual staining applications. It guides the training of the generator towards generating virtually-stained images that resemble authentic images in terms of perceptual content.

The first two networks have been described in detail in a previous work [29]. The additional perceptual discriminator consists of a Densely Connected Convolutional Networks (DenseNet121) [30], pre-trained on the ImageNet dataset. This network receives virtually- and chemically-stained images and maps them into an N-dimensional representation to assess their similarities in feature space [31]. This approach allows the network to evaluate discrepancies in content and style, improving the generator’s performance to reproduce cell elongations and internal texture and discern cell boundaries in high-density scenarios. Importantly, this network remains fixed during the cGAN training, and no changes are applied to its pre-trained weights.

To obtain quantitative information about the sample’s viability we used a deep-learning method to train neural networks using only a single label-free image to track particles called LodeSTAR.[14] This method forms part of the Python library DeepTrack 2.1.[29] We based our counting networks on one of the provided example notebooks[32] and trained four separate networks, one for each dye (live or apoptotic) and type of data (chemically- or virtually-stained) by choosing representative images for these four cases.

Time evolution of the number of live or apoptotic cells from the internal validation dataset at low flow velocity when stained chemically or virtually is represented in Figure 2a. We acquired the data for 12 hours, but cells exposed to calcein suddenly ruptured after approximately half of the experiment. Therefore, we chose images for both calcein and caspase internal validation within the first five hours. All data were normalised to an average of the first three data points for each dye and type. The number of live cells is stable over time in the virtually-stained images but decays to almost half in 5 hours in the chemically-stained ones. The discrepancy between the number of chemically- and virtually-stained viable cells can be explained by several mechanisms: First, in our experimental approach, the chemical stain is mixed with the medium and continuously flows through the cell culture. There are two consequences of this. (i) The administrated calcein concentration is higher than recommended by the manufacturer, and self-quenching could occur. (ii) The cells’ constant exposure to dye and illumination light for hours becomes cytotoxic (what we observed in the internal validation data set). Either the mechanism, the calcein fluorescence intensity in affected cells decreases (meanwhile the background increases) and subsequently the cells are ignored by the automatic counting network. This is not seen in the virtually stained images as the phase contrast images remain unaffected. The cytotoxic effect (see Movies S1-S2) was confirmed in a control experiment where cells were only subjected to culture medium and imaged with phase-contrast microscopy (see Movie S3). The control experiment shows that cells keep their morphological phenotype throughout the experiment compared to the calcein-exposed cells.

**FIG. 2.**
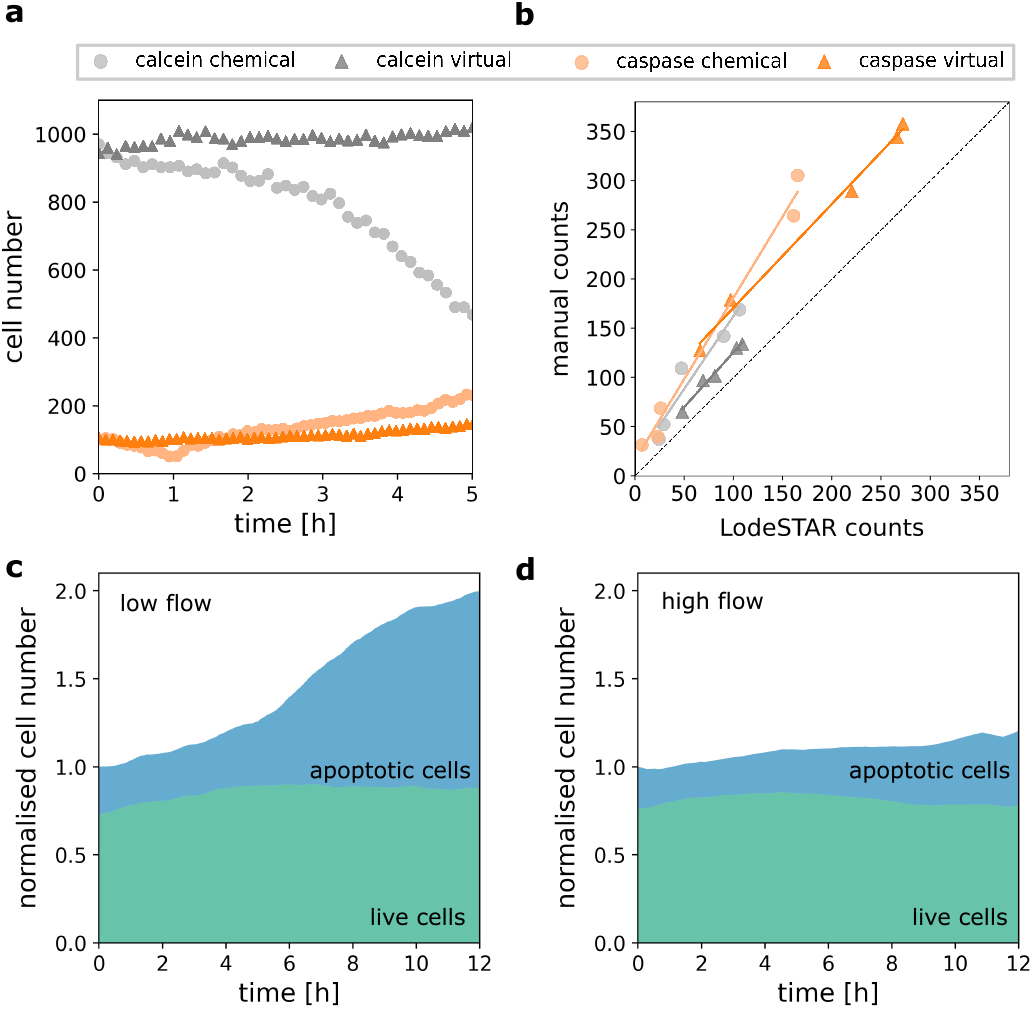
Evaluation of the virtual staining and automatic cell counting. (a) Time evolution of the number of live and apoptotic cells when stained chemically or virtually (colour-coded) during two different low flow experiments. The displayed data were normalised to an average of the first three data points for each dye and type. (b) Comparison of manual counting versus LodeSTAR network output. Five images of each type were manually-counted by two independent counters and the average of those counts was compared with the automatic counting approach. (c-d) The trained LodeSTAR networks were used to analyse the virtually-stained phase-contrast images of cells cultured under low (c) and high (d) flows. The data were normalised to the total number of cells at time zero for each experiment and averaged by using a rolling average with window size 5.

For apoptotic cells, the trend is similar for the number of cells in chemically- and virtually-stained images. There are slightly fewer cells counted as apoptotic in the virtually-stained images than in the chemically-stained ones.

Figure 2b shows the correlation between manual and automatic counting with LodeSTAR. The number of cells in twenty images, five per dye and staining type, was manually counted by two independent counters. The plot represents the correlation between an average of the results of the two counters and the automatic counting done by LodeSTAR. The human eye generally tends to classify more spots as cells than the neural network. As mentioned previously, the low-intensity caspase signal in healthy cells might result in subjective analysis. We considered this when training the LodeSTAR, resulting in a slightly more restrictive network than manual counters.

In Figure 2c-d, the time evolution of cells automatically stained by the cGAN as live or apoptotic when cultured in low (c) or high (d) flow and subsequently counted by LodeSTAR is shown. The presented data is the result of virtual staining of the phase-contrast images from the experiments with chemical caspase-3/7 dye as experiments with calcein suffered from the problems mentioned previously. For visualisation, the data were averaged using a rolling average with a window size of 5 data points and normalised to the total number of cells (live+apoptotic) at the time zero. Over time, the number of live cells is stable regardless of the flow rate exposure. These results support our previous reasoning that the virtually-stained cells are not affected by the self-quenching mechanism.

The number of apoptotic cells increases in both flow experiments over time but at different rates. Cells exposed to low flow suddenly turn apoptotic after approximately 5 hours into the experiment, compared to the cells cultured under high flow. This is expected as the cells cultured in low flow are exposed to a lower rate of media replenishment, which is detrimental to the cells in the long run. Cells cultured under high flow also undergo apoptosis but at a much slower rate. Again, this is possibly caused by the continuous exposure to illumination light and chemical dyes.

Apoptosis is an active process and requires cells to be alive, i.e., having an intact membrane. The total number of cells is seemingly increasing over the time course of the experiment, a behaviour more evident for the low-than the high-flow cell culture condition. This mirrors the used dyes’ working principles, where most apoptotic cells are stained positive for both calcein and caspase.

To conclude, we have developed a non-invasive method based on deep learning to discriminate between alive HMEC-1 cells and cells that have initiated apoptosis by following their viability over time. This approach is based on virtually-stained images obtained from time-lapse phase-contrast images of the sample and further cell counting to attain live versus apoptotic cell ratios for different experimental conditions. These microscopy images provide enough information about the thin HMEC-1 cells for the network to distinguish between live and apoptotic cells based on their morphology and possibly internal subcellular structures. In physiologically-relevant studies, cells have cell–cell contact, and corresponding in vitro experiments result in high confluency. Even if optimal for the cellular function, the resulting non-homogeneous 2D cell layer is difficult to analyse and, specifically, to stain with standard chemical dyes. The microscopy imaging and subsequent cell counting using traditional approaches are also very challenging and may force investigators to dilute the cell sample even to perform the analysis. A lower magnification speeds up the monitoring and enables capturing dynamic cell changes in large cell populations. Once again, while optimal for high throughput and statistics, the large amount of data to process restricts manual cell counting and urges for automatic analysis.

Our presented approach is much less labour-intensive and cell-invasive than standard fluorescent stain-dependent methods. It may be used for further functional studies on HMEC-1 cells or other cells that thrive in high perfusion environments where a continuous infusion of a chemical dye is required to follow long-term dynamic events. Our cell viability method allows for studying other processes that still require fluorescent probes and chemical staining. Consequently, it facilitates a higher information gain on the same cell population than relying only on the conventional chemical staining procedures.

## FUNDING AND ACKNOWLEDGMENTS

This project has received funding from the European Union’s Horizon 2020 research and innovation programme under the Marie Sklodowska-Curie grant agreement No. 766181.

## AUTHOR CONTRIBUTIONS

Author contributions are defined based on the CRediT (Contributor Roles Taxonomy) and listed alphabetically. Conceptualization: C.B.A., S.H., Z.K. Formal analysis: S.H., Z.K., J.P. Funding aquisition: C.B.A., M.G., Investigation: Z.K. Methodology: S.H., Z.K., J.P. Project administration: C.B.A., G.V. Software: S.H., Z.K., J.P. Supervision: C.B.A. Validation: C.B.A, S.H., Z.K., J.P. Visualization: Z.K., J.P. Writing – original draft: C.B.A., Z.K., J.P. Writing – review & editing: C.B.A, S.H., Z.K., J.P., G.V.

## COMPETING INTERESTS STATEMENT

The authors declare no competing interests.

## DATA AVAILABILITY

Examples of data and software that support the findings of this study are openly available on our Github repository https://github.com/softmatterlab/live-apoptotic-virtual-staining.

## Notes

https://github.com/softmatterlab/live-apoptotic-virtual-staining

### Competing Interest Statement

The authors have declared no competing interest.

