## Supplementary Information for "Dynamic live/apoptotic cell assay using phase-contrast imaging and deep learning"

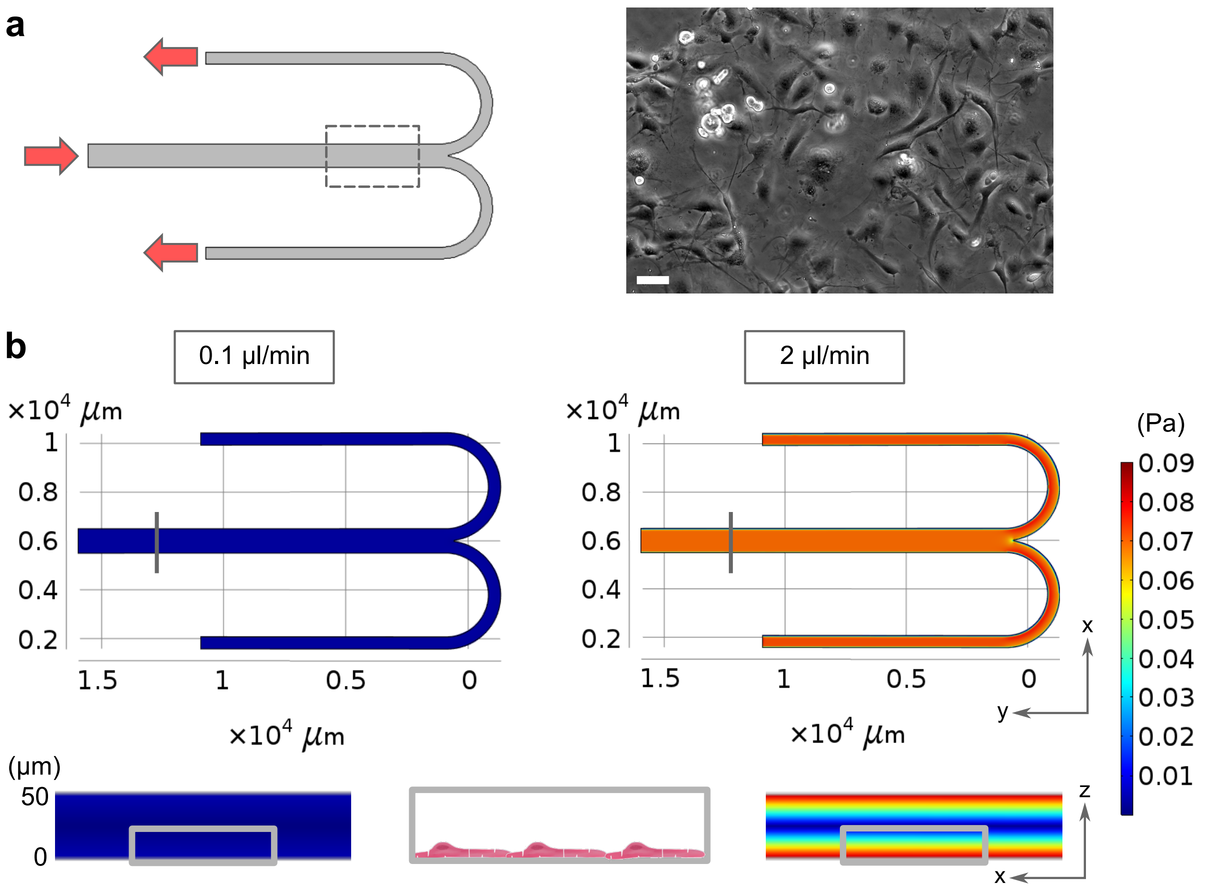


FIG. S1. **Microfluidic device design and shear stress simulations.** **a** Schematic representation of experimental conditions. Cells are introduced into the centre channel, where they are allowed to sediment before the media flow is started (flow direction according to the arrows). The experiment is performed under two different flows, low (0.1 µl/min) and high (2 µl/min), to generate ground truth data for the neural network. Phase-contrast image of HMEC-1 cells is shown in the inset on the right. Scale bar is 50 µm. **b** Device dimensions and 3D shear stress COMSOL Multiphysics simulations for the two flows used in the experiments. Centre channel cross-sections are presented in the lateral insets displaying the shear stress distribution inside the channel. In the case of low flow, the differences across the channel are negligible, but this is not the case for high flow. Because endothelial cells are very thin (below 10 µm), the HMEC-1 cells are exposed to 4 mPa and 80 mPa shear stress values for low and high flow, respectively, as shown in the centre inset.

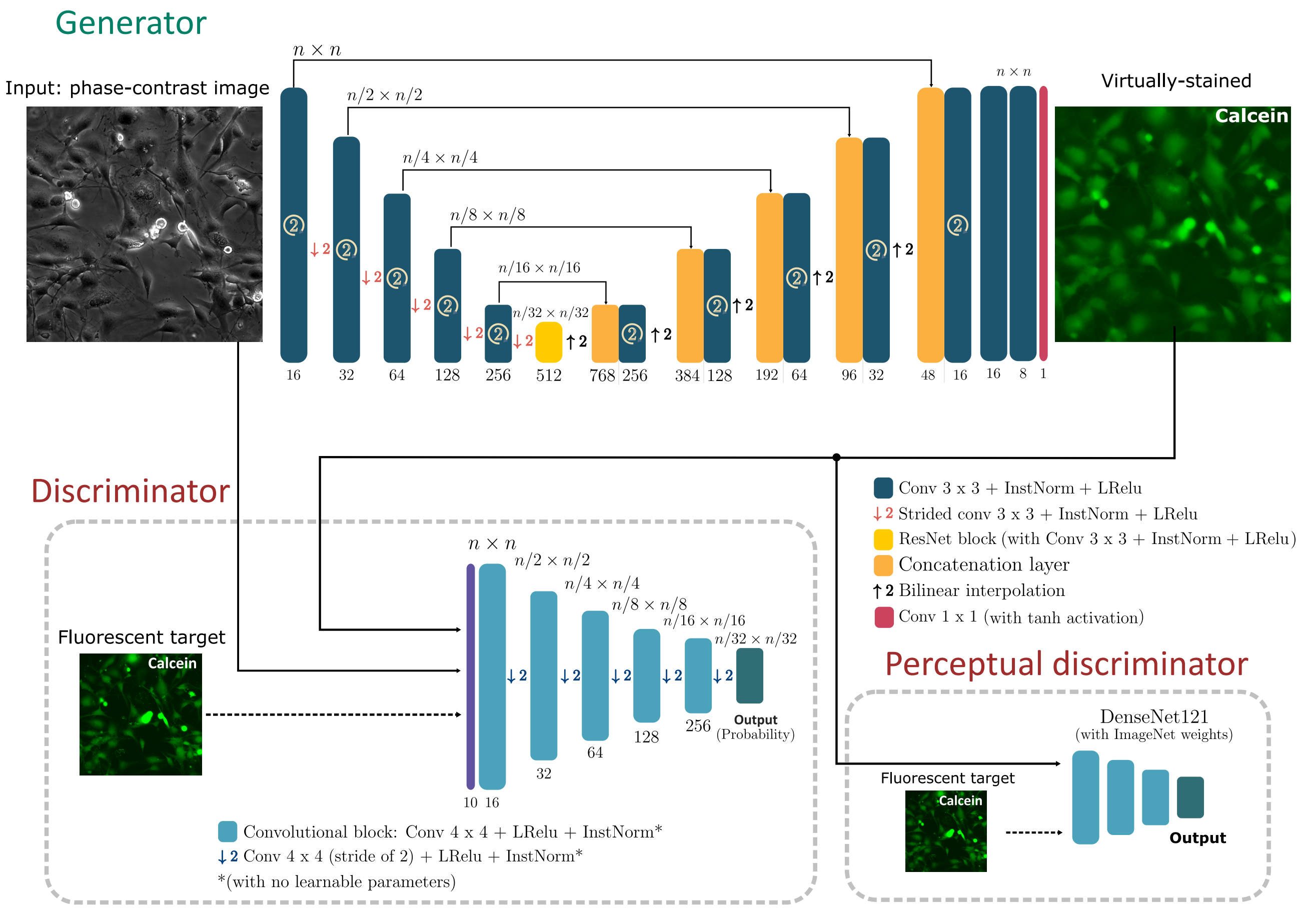


FIG. S2. **The architecture of the cGAN used for virtual staining.** The generator transforms an input phase-contrast image into an image of virtually-stained cells as live or apoptotic (there are two separate cGAN networks, one for each type of dye). It follows a U-Net architecture, with symmetric encoder and decoder paths connected through skip connections. The encoder is coupled to the decoder through a residual network (ResNet) block with 512 feature channels. The U-Net decoder uses bilinear interpolation for upsampling. The upsampled features are concatenated with the corresponding feature map and processed by convolutional layers. The generator uses instance normalisation and leaky ReLU activation for all layers except for the final convolutional layer and the upsampling layers. The discriminator is a PatchGan that receives both the phase-contrast and fluorescence images, which can be either chemically- or virtually-stained, and divides them into overlapping patches that are analysed and labelled as real or fake. The network architecture comprises $4 \times 4$ convolutional layers followed by strided convolutions for downsampling. All layers use instance normalisation (with no learnable parameters) and leaky ReLU activation. The perceptual discriminator is a DenseNet121 pretrained on the ImageNet dataset. It receives images of both chemically- and virtually-stained cells and maps them into an N-dimensional representation to assess their similarities in feature space. This network remains fixed during the cGAN training, and no changes are applied to its pretrained weights. The same approach was used to train the cGAN for caspase-3/7 virtual staining.

**Supplementary Movie S1** Phase-contrast images from an experiment monitoring endothelial cells exposed to high flow and calcein AM. The presented time-lapse images (from T=0 min to T=110 min) reveal the cytotoxicity of the used dye. Initially cells maintain their normal elongated morphology. However, one cell (in the upper right corner) rounds up and eventually bursts, forming typical apoptotic bodies (small droplet-shaped structures).

**Supplementary Movie S2** Fluorescence images from an experiment monitoring endothelial cells exposed to high flow and calcein AM. The presented time-lapse images (from T=0 min to T=110 min) reveal that all cells are stained positive with calcein. The calcein intensity increases as the cell morphology changes (as seen in the representative cell in the upper right corner). By the movie's end, the calcein intensity in this cell is very high, reaching its highest value just before the cell ruptures. This video corresponds to the fluorescence modality of the same experiments as movie S1.

**Supplementary Movie S3** Endothelial cells monitored with phase-contrast microscopy in a high flow environment display healthy morphology. Phase-contrast microscopy time-lapse images (from T=0 min to T=110 min) of endothelial cells exposed to only medium reveal the imaged cells' normal, elongated morphology throughout the movie's time frame.
